# “METAVIR score Ultrasound” – Pictorial Essay of Anatomy / Ultrasound / Elastography / Histology

**DOI:** 10.1101/2022.08.06.503057

**Authors:** Luís Jesuino de Oliveira Andrade, Luís Matos de Oliveira, Gabriela Correia Matos de Oliveira

## Abstract

The METAVIR score was specifically elaborated and evaluated in chronic hepatitis C, is scoring system for evaluation the inflammatory activity and fibrosis on liver biopsies, although it ultrasound may indicate the fibrosis degree.

**Objective:** To present a pictorial essay by composing histological, ultrasound and elastography images of the METAVIR score of fibrosis.

**Method:** Using software for image composition, based on the sonographic image of the liver with degrees of fibrosis, secondary to hepatitis C, we adapted the corresponding histological image (METAVIR) and presented a pictorial essay.

**Results:** The correlation of the anatomical image with the sonographic image and of the histological image with the corresponding sonographic image and elastography image are demonstrated.

**Conclusion:** Ultrasonographic features and elastography in evaluation hepatic fibrosis present in hepatitis C can correlate with histological features METAVIR score.

## Introduction

The hepatitis c virus was discovered in 1989,^1^ and since its discovery several studies have been elaborated to evaluate its interaction with the hepatocyte.^2^ The World Health Organization has set a goal of eliminating hepatitis C by 2030, a condition defined as a 65% reduction in mortality from HCV and a 90% reduction in new infections.^3^

Liver biopsy is considered the gold standard for fibrosis assessment, and the degree of fibrosis and inflammation is classified by the METAVIR score system a semi-quantitative system that was primarily developed and studied in individuals with chronic hepatitis C. The METAVIR score is a system consisting of a coding form of two letters and two numbers where A encodes the degree of histological activity and F encodes the degree of fibrosis. Thus, A0: no activity, A1: mild activity, A2: moderate activity, and A3: severe activity, while F0: no fibrosis; F1: portal fibrosis without septa, F2: portal fibrosis with few septa, F3: numerous septa without cirrhosis, and F4: cirrhosis.^4^

The assessment of liver stiffness is of fundamental importance in the treatment of liver diseases because the prognosis of the disease is directly linked to the progression of liver fibrosis.

Ultrasound is a low-cost method, easy to perform, uses no ionizing radiation, is well accepted by patients. Ultrasonography is an essential diagnostic imaging method in clinical practice for evaluation of the liver. Sonographic studies have demonstrated a strong correlation between histological and echography liver fibrosis findings.^5^ Ultrasonographic findings that include irregularity or rippling of the liver surface, coarse echotexture, and focal nodulations of the liver parenchyma, and diffuse periportal thickening are correlated with liver fibrosis at various stages.^6^

Elastography is a developed in the 1990’s non-invasive imaging technique based on the tissue mechanical properties. It is an alternative method to liver biopsy in which stiffness correlated to fibrosis of the liver parenchyma are measured, and its effectiveness has been reported in diagnosing the degree of liver fibrosis.^7^ The currently available ultrasound elastography techniques are strain imaging and shear wave imaging. The strain imaging technique consists of applying an external stress to the liver tissue which provides a qualitative assessment, while the shear wave imaging technique a dynamic stress is applied to the liver tissue using a vibrating mechanical device.^8^

Despite the various sophisticated imaging examinations used today, numerous misinterpretations of images can occur. The purpose of this study is to present a pictorial essay by composing histological, ultrasound and elastography images of the Metavir score of fibrosis.

## Method

Using software for image composition, paint one of the basic windows programs, based on the ultrasound image of the liver with the various degrees of fibrosis and correlating it with the corresponding histological image (METAVIR score) we present a pictorial essay.

Correlation of ultrasound images, with interest in the degree of fibrosis by sonographic signs of liver fibrosis, and review of histological specimens of the various degrees of liver fibrosis classified by the METAVIR score were performed by an ultrasound specialist and a pathologist respectively.

This study proposes to investigate the theoretical understanding of pathologies of clinical practice, and according to the Research Ethics Committee of Brazil (CEP), CNS Resolution 510/2016, there is no need for CEP evaluation.

## Resuts and Discussion

### METAVIR score F0

#### Anatomy

The liver is an organ located in the right hypochondrium in direct contact with the gallbladder, stomach and diaphragm, is covered by Glisson’s capsule, and is almost entirely covered by the visceral peritoneum in association with peritoneal ligaments.^9^

#### Ultrasonography

The sonographic image of normal liver parenchyma has little variation between individuals. The echotexture of the normal liver parenchyma has a homogeneous mean echogenicity, slightly less echogenic than the spleen and slightly brighter than the renal cortex when compared at the same depth level.^10^

#### Elastography

Based on several studies and based on the so-called ‘rule of four’ (5, 9, 13, and 17 kPa) for acoustic radiation force impulse (ARFI) techniques used to evaluate liver fibrosis of viral etiology liver stiffness of 5 kPa (1.3 m/sec) or less has a high probability of being normal.^11^

#### Histology

The liver tissue histologically is composed of hexagon-shaped lobules, and within the lobules the hepatocytes are aligned as cellular strands connecting the peripheral portal tracts with the central veins. The hepatic sinusoid which corresponds to a specialized capillary system is traversed by the hepatic artery and portal vein. The combination of the hepatic canaliculi will form the bile ducts that associated with the hepatic artery and a branch of the portal vein will constitute the “portal triad”.^12^

Figure 1 shows the composition of images that correspond to the normal liver (anatomy, ultrasound, elastography, and histology).

**Figure 1.**
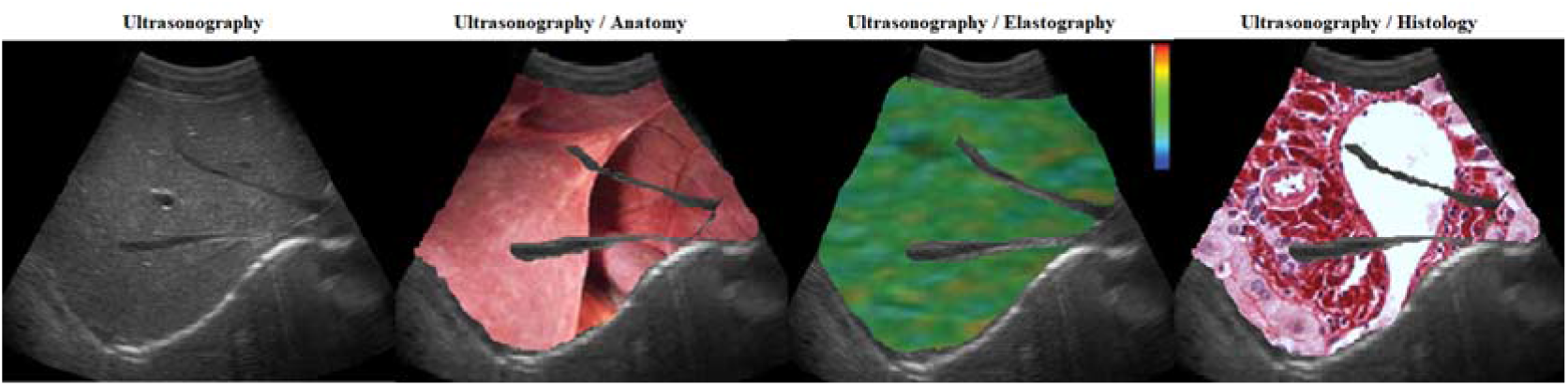
Normal Liver. **Source**: Research result

### METAVIR score F1

#### Anatomy

From an anatomical point of view, a fibrosis score of F1 from the METAVIR score shows no significant anatomical changes when compared to a liver without fibrosis.

#### Ultrasonography

Echographically the F1 score of the METAVIR score is characterized by mildly increased echogenicity of the walls of the main portal vein and reduced caliber of the portal vessels, and diffuse linear echogenicity resulting from mild diffuse periportal thickening and diffuse echogenicity in the hepatic periphery may also be found.^13^

#### Elastography

The ARFI technique in the assessment of fibrosis score, where there is fibrosis with portal zone expansion with hepatic stiffness to elastography in the range of 5.3-7.4 kPa with a mean of 6.43 kPa (1.47 m/s) corresponds to METAVIR score F1.^14^

#### Histology

The F1 stage of the METAVIR score presents as anatomopathological characteristics portal fibrosis without septa corresponding to incipient fibrosis or minimal liver fibrosis.^4^

Figure 2 shows the composition of images that correspond to the METAVIR score F1 liver (anatomy, ultrasound, elastography, and histology).

**Figure 2.**
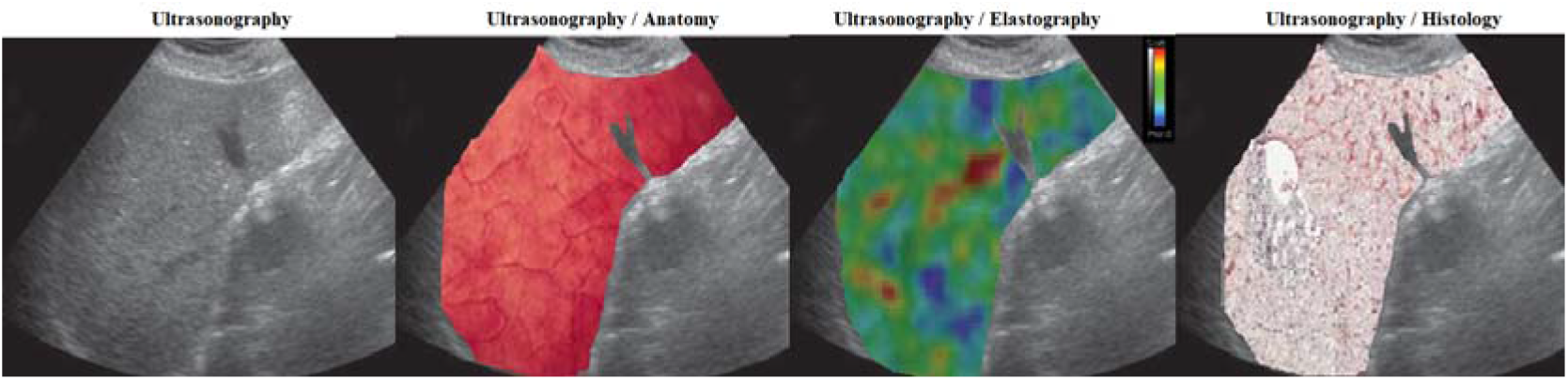
METAVIR score F1 **Source**: Research result

### METAVIR score F2

#### Anatomy

Macroscopic anatomy of the liver shows a granular appearing surface (regenerative nodules), presence of fibrous tissue septa that subdivide the liver in a pattern similar to a puzzle.

#### Ultrasonography

Slight echogenic thickening of the walls of two or more peripheral portal branches, with little or no thickening of the wall of the main portal vein and with slight narrowing of the portal radicles. There is wall thickening of the gallbladder. In this grade, periportal fibrosis is much more evident than in the previous grade, and irregular narrowing of the lumen of the portal vessels is noted.^15^

#### Elastography

In the diagnosis of stage F2 liver fibrosis, the liver stiffness meters for elastograph liver fibrosis, where fibrosis occurs with expansion in most portal zones and occasional fibrosis with signaling fiber bridges, if by the ARFI technique in the evaluation of liver stiffness is in the range between 7.1 kPa (1.59 m/s) corresponding to the METAVIR F2 score.^16^

#### Histology

The F2 stage of the METAVIR score presents as anatomopathological characteristics portal fibrosis with rare septa extending into the lobules, corresponding moderate fibrosis.^4^

Figure 3 shows the composition of images that correspond to the METAVIR score F2 liver (anatomy, ultrasound, elastography, and histology).

**Figure 3.**
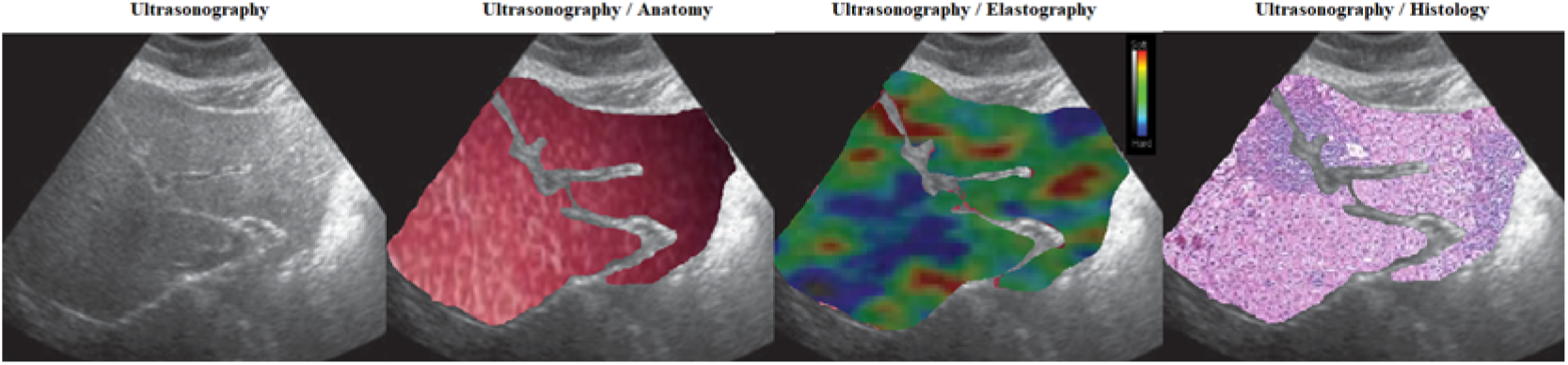
METAVIR score F2 **Source**: Research result

### METAVIR score F3

#### Anatomy

The liver presents with a pronounced nodularity on the external surface and a shiny capsule. In addition, the organ presents with reduced volume and a hardened consistency.

#### Ultrasonography

Echographically, servera fibrosis is characterized by severe, irregularly distributed periportal thickening of most portal radicles with marked narrowing of the central sonolus. The thickening is marked at the bifurcation of the portal vein, extending to the liver surface. The wall of the main portal vein is thickened by 2 to 10 mm, as is the wall of the gallbladder.^17^

#### Elastography

Using the ARFI technique to assess the fibrosis score, where liver fibrosis characterized by numerous septa, elastography shows an elasticity quantification of 9.5 kPa and a shear wave velocity of 1.74 m/s which corresponds to the METAVIR F3 score.^18^

#### Histology

The F3 stage of the METAVIR score presents as anatomopathological characteristics numerous septa extending into the adjacent portal tracts or terminal hepatic venules, corresponding severe fibrosis.^4^

Figure 4 shows the composition of images that correspond to the METAVIR score F3 liver (anatomy, ultrasound, elastography, and histology).

**Figure 4.**
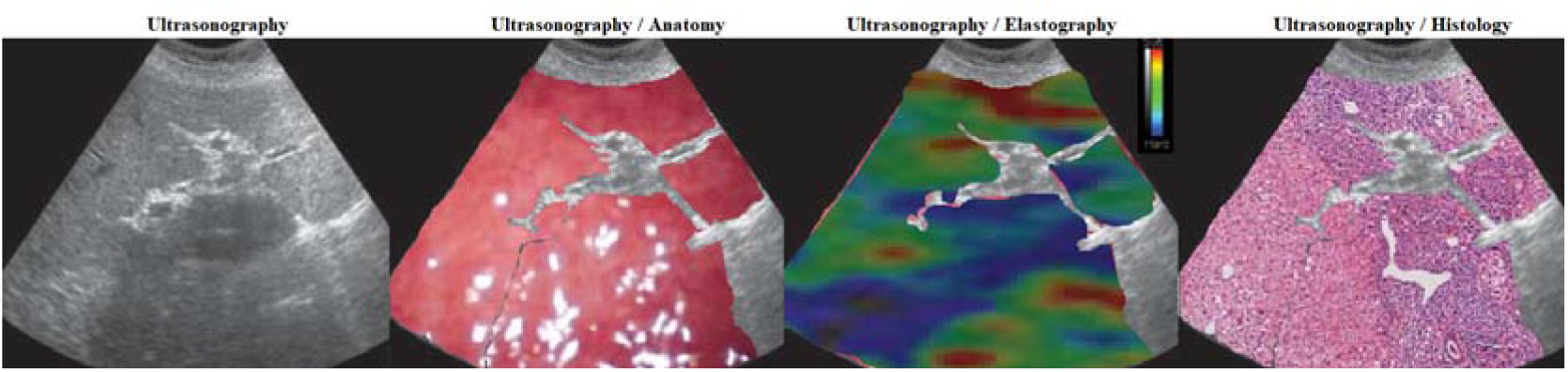
METAVIR score F3 **Source**: Research result

### METAVIR score F4

#### Anatomy

The external surface of the liver presents with pronounced nodularity and a shiny capsule. Organ with important reduction in volume, hardened consistency, pale to brownish color.

#### Ultrasonography

Ultrasonography is a well-established non-invasive imaging test for the diagnosis of liver cirrhosis. The echographic alterations are due to the progression of hepatic fibrosis that will be characterized by an increase in the volume of the liver, echographic coarse or nodular texture of the hepatic parenchyma and, with the evolution of portal hypertension, an increase in the caliber of the portal vein and flow inversion of this vein, visualized with the aid of Doppler. Thus, ultrasonography is a low-cost and easily accessible imaging technique to diagnose cirrhosis.^19^

#### Elastography

The use of elastography has been used to constantly improve the diagnosis of cirrhosis. Thus, liver cirrhosis on elastography evaluation using the ARFI technique defined an elasticity quantification of 12.5 kPa and a shear wave velocity of 1.92cm corresponding to the METAVIR F4 score.^20^

#### Histology

Cirrhosis according to the Laennec histological classification correlates with different stages of clinical severity. The staging of cirrhosis according to the histological features described by Laennec was modified by the METAVIR system by subdividing the highest degree of fibrosis (F4) into 4A, 4B and 4C in order to differentiate the changes in the severity of cirrhosis.^21^

Figure 5 shows the composition of images that correspond to the METAVIR score F4 liver (anatomy, ultrasound, elastography, and histology).

**Figure 5.**
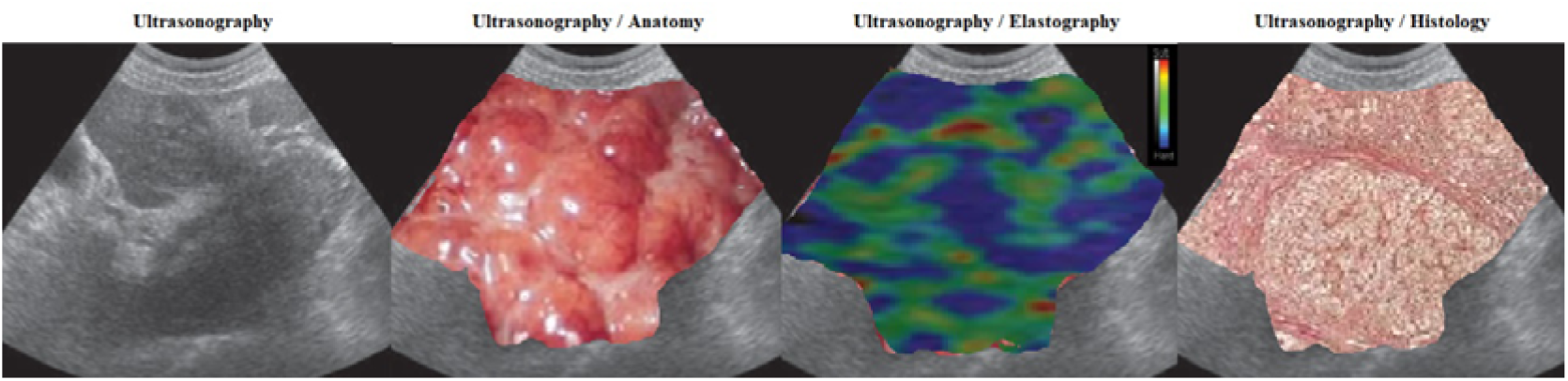
METAVIR score F4 **Source**: Research result

## Conclusion

Ultrasonographic features and elastography in evaluation hepatic fibrosis present in hepatitis C can correlate with histological features METAVIR score.

## References

1. Choo QL, Kuo G, Weiner AJ, Overby LR, Bradley DW, Houghton M. Isolation of a cDNA clone derived from a blood-borne non-A, non-B viral hepatitis genome. Science. 1989;244(4902):359–62.

2. Zeisel MB, Felmlee DJ, Baumert TF. Hepatitis C virus entry. Curr Top Microbiol Immunol. 2013;369:87–112.

3. World Health Organization Global Health Sector Strategy on Viral Hepatitis 2016–2021. Towards Ending Viral Hepatitis. [(accessed on July 28,2022]. Available online: https://apps.who.int/iris/bitstream/handle/10665/246177/WHO-HIV-2016.06-eng.pdf [Ref list].

4. Bedossa P, Poynard T. An algorithm for the grading of activity in chronic hepatitis C. The METAVIR Cooperative Study Group. Hepatology. 1996;24(2):289–93.

5. Nishiura T, Watanabe H, Yano K, Ito M, Abiru S, Fujimoto T, et al. Integrated fibrosis scoring by ultrasonography predicts the occurrence of hepatocellular carcinoma in patients with chronic hepatitis C virus infection. J Med Ultrason (2001). 2011;38(1):13–9.

6. Zheng J, Guo H, Zeng J, Huang Z, Zheng B, Ren J, et al. Two-dimensional shear-wave elastography and conventional US: the optimal evaluation of liver fibrosis and cirrhosis. Radiology. 2015;275(1):290–300.

7. Agbim U, Asrani SK. Non-invasive assessment of liver fibrosis and prognosis: an update on serum and elastography markers. Expert Rev Gastroenterol Hepatol. 2019;13(4):361–74.

8. Ozturk A, Grajo JR, Dhyani M, Anthony BW, Samir AE. Principles of ultrasound elastography. Abdom Radiol (NY). 2018;43(4):773–85.

9. Skandalakis JE, Skandalakis LJ, Skandalakis PN, Mirilas P. Hepatic surgical anatomy. Surg Clin North Am. 2004;84(2):413–35,

10. Khan SA, Yasmeen S, Adel H, Adil SO, Huda F, Khan S. Sonographic Evaluation of Normal Liver, Spleen, and Renal Parameters in Adult Population: A Multicenter Study. J Coll Physicians Surg Pak. 2018;28(11):834–39.

11. Barr RG, Ferraioli G, Palmeri ML, Goodman ZD, Garcia-Tsao G, Rubin J, et al. Elastography Assessment of Liver Fibrosis: Society of Radiologists in Ultrasound Consensus Conference Statement. Radiology. 2015;276(3):845–61.

12. Gerber MA, Thung SN. Histology of the liver. Am J Surg Pathol. 1987;11(9):709–22.

13. Li J, Qureshi M, Gupta A, Anderson SW, Soto J, Li B. Quantification of Degree of Liver Fibrosis Using Fibrosis Area Fraction Based on Statistical Chi-Square Analysis of Heterogeneity of Liver Tissue Texture on Routine Ultrasound Images. Acad Radiol. 2019;26(8):1001–1007.

14. Fraquelli M, Baccarin A, Casazza G, Conti CB, Giunta M, Massironi S, et al. Liver stiffness measurement reliability and main determinants of point shear-wave elastography in patients with chronic liver disease. Aliment Pharmacol Ther. 2016;44(4):356–65.

15. Zheng RQ, Wang QH, Lu MD, Xie SB, Ren J, Su ZZ, et al. Liver fibrosis in chronic viral hepatitis: an ultrasonographic study. World J Gastroenterol. 2003;9(11):2484–9.

16. Castéra L, Vergniol J, Foucher J, Le Bail B, Chanteloup E, Haaser M, et al. Prospective comparison of transient elastography, Fibrotest, APRI, and liver biopsy for the assessment of fibrosis in chronic hepatitis C. Gastroenterology. 2005;128(2):343–50.

17. Wakui N, Nagai H, Yoshimine N, Amanuma M, Kobayashi K, Ogino Y, et al. Flash Imaging Used in the Post-vascular Phase of Contrast-Enhanced Ultrasonography is Useful for Assessing the Progression in Patients with Hepatitis C Virus-Related Liver Disease. Ultrasound Med Biol. 2019;45(7):1654–62.

18. Talwalkar JA, Kurtz DM, Schoenleber SJ, West CP, Montori VM. Ultrasound-based transient elastography for the detection of hepatic fibrosis: systematic review and meta-analysis. Clin Gastroenterol Hepatol. 2007;5(10):1214–20.

19. Sharma S, Khalili K, Nguyen GC. Non-invasive diagnosis of advanced fibrosis and cirrhosis. World J Gastroenterol. 2014;20(45):16820–30.

20. Ziol M, Handra-Luca A, Kettaneh A, Christidis C, Mal F, Kazemi F, et al. Noninvasive assessment of liver fibrosis by measurement of stiffness in patients with chronic hepatitis C. Hepatology. 2005;41(1):48–54.

21. Rastogi A, Maiwall R, Bihari C, Ahuja A, Kumar A, Singh T, et al. Cirrhosis histology and Laennec staging system correlate with high portal pressure. Histopathology. 2013;62(5):731–41.

